# Intra-arterial Deoxyribonuclease therapy improves stroke outcomes in aged mice

**DOI:** 10.1101/2024.10.28.620095

**Authors:** Junxiang Yin, Michael Wu, Jennifer White, Ellie StClair, Michael F. Waters

## Abstract

**Background:** Futile recanalization affects more than half of acute ischemic stroke (AIS) patients. Neutrophil extracellular traps (NETs) are a major factor of microvascular hypoperfusion after stroke. Deoxyribonuclease I (DNase) targeting NETs exhibited a neuroprotective effect in young mice with AIS. This study explored a novel direct intra-arterial administration of DNase therapy and its effect in aged mice with AIS.

**Method:** AIS was induced in aged C57BL/6 mice followed by reperfusion and immediate, intra-arterial DNase administration via the internal carotid artery. Cerebral blood flow, neurological function, cerebral infarct volume, and NET markers were examined.

**Results:** Direct intra-arterial DNase therapy significantly increased cerebral blood flow, reduced neurological deficit scores, increased the latency to fall in wire hang test, reduced cerebral infarct volume, and decreased neutrophil and NET count in both the parenchyma and micro vessels in aged mice with AIS compared with age-matched, vehicle controls.

**Conclusion:** Our data is the first to demonstrate that successful, direct intra-arterial DNase therapy provides more efficient cerebral reperfusion and better outcomes after recanalization during the treatment of large vessel occlusion in aged mice. This study provides evidence for the potential clinical application of catheter delivered intra-arterial DNase therapy post-recanalization.

## Introduction

Stroke is a leading cause of mortality and disability worldwide [1, 2]. More than 30% of acute ischemic strokes (AIS) are large vessel occlusion (LVO), disproportionately contributing to 95% of mortalities and 62% of long-term disability [3]. Endovascular thrombectomy (EVT) is the standard of care for LVOs [4-6]. Despite there being high success rates of recanalization with EVT, more than half of patients still suffer poor outcomes and low micro-perfusion, a phenomenon coined futile recanalization [7-10]. This futile recanalization is recognized as a challenge during AIS treatment and determining the mechanisms are integral to improving the outcomes of stroke patients [11-14]. Numerous factors are associated with the failure to restore penumbral microcirculation, one of which is microvascular thrombosis resulting in distal capillary hypoperfusion.

Neutrophil Extracellular Traps (NETs) are extracellular deoxyribonucleic acid (DNA) networks wrapped around histones and granular proteins produced and extruded by activated neutrophils. Neutrophils and NETs are important components of cerebral thrombi [15] and markers of NET formation are also significantly increased in the plasma and cerebral penumbra of ischemic stroke patients [16, 17]. Both neutrophils and NETs are elevated in the central nervous system and in the peripheral circulation of ischemic stroke patients and animal models of ischemic stroke. Elevated plasma NET biomarkers are also correlated with worse stroke outcomes [16-19]. Additionally, NETs have been identified as major triggers and structural factors of microvascular thrombosis, and they exacerbate microvascular hypoperfusion in stroke [17, 20, 21]. Furthermore, NETs are thought to play a key mechanistic role in thrombolysis resistance [22]. Recent studies have also reported that an early decline in deoxyribonuclease (DNase) activity and elevated NET marker concentrations in circulation increased the risk of delayed cerebral ischemia following hemorrhagic stroke [23, 24]. By cleaving DNA, DNase has been proposed as an efficient antithrombotic drug [25]. Additionally, DNase has exhibited promising neuroprotective effects in acute ischemic stroke via targeting NETs in young mouse models of ischemic stroke [15, 18, 19]. Intraperitoneal injection (IP) and intravenous tail vein injection (IV) injection are two of the most common administration routes in small animal experiments. However, IP injection is unlikely to be clinically translatable. Tail vein injection is a systemic administration requiring a high dose and potential repeated administrations limits this application. To date, there are no FDA approved therapies for modifying NETs in stroke. To explore a more efficient, clinically translatable, and low dose strategy for DNase therapy and its effects during the treatment of AIS, we investigated a novel, direct intra-arterial DNase therapy in aged mice with AIS.

## Materials and Methods

### Animals

Aged C57BL/6 mice (∼20 months) were used in this study. All experiments involving animals were performed following the protocols in accordance with the Revised Guide for the Care and Use of Laboratory Animals and were approved by the Institutional Animal Care and Use Committee (IACUC) of the Icahn School of Medicine at Mount Sinai. The study was carried out in compliance with the ARRIVE guidelines. All methods were performed in accordance with the relevant guidelines and regulations.

### Deoxyribonuclease I (DNase) activity assessment *in vitro* and *in vivo*

DNase activity *in vitro* test: Recombinant DNase I (DNase, Sigma-#4536282001) was diluted with pure water into different concentrations (0.01, 0.001, 0.0001, 0.00001 ug/µl); heat-inactivated DNase was used as a control (HI-DNase, 0.01 ug/µl). DNase activity was measured with a DNase activity fluorescent assay kit (Acrobiosystems, #ASE-1002). Relative fluorescence units (RFU) were used for DNase activity curve analysis. DNase activity *in vivo* test: 6-month-old wild type mice received DNase (6mg/kg, 1 Unit DNase = 1 ug DNase) via intraperitoneal (IP) injection (n=3) or tail vein (IV) injection (n=4). Age-matched wild type mice receiving a vehicle injection were used as controls (Ctrl, n=3). Serum was collected 30 minutes after DNase or vehicle administration and used to evaluate DNase activity.

### Transient middle cerebral artery occlusion (MCAO)

Mice were subjected to the intraluminal filament method of transient middle cerebral artery occlusion (MCAO) as described previously [26]. Briefly, mice were anesthetized and placed in the supine position on a heating pad, and body temperature was monitored using the small animal anesthesia system (Kent Scientific Cor. FL). Under a stereo dissecting microscope (AmScope, CA), the right common carotid artery (CCA), external carotid artery (ECA), and internal carotid artery (ICA) were exposed and isolated. To induce MCAO, a silicon-coated nylon monofilament (Doccol, MA) was introduced into the internal carotid artery through the external carotid artery, then further advanced to the origin of the MCA. Successful MCAO was confirmed by ≥75% decrease in cerebral blood flow (CBF) monitored by laser doppler flowmeter (MoorVMS-LDF2, Moor Instruments, UK). After 60 minutes of ischemia, the filament was fully withdrawn for reperfusion. Buprenorphine SR was administered at 0.05 mg/kg of body weight to alleviate any postoperative pain, and the mice were put into a portable animal intensive care unit to recover before being transferred to their home cages.

### Evaluation of Cerebral blood flow

CBF was evaluated using 2D laser speckle system (PeriCam PSI-Z, Perimed). Mice were anesthetized and laid prone on a pre-warmed SurgiSuite surgical platform (Kent Scientific, CT). A small incision was made, and the skin was retracted to expose the intact skull. Cerebral perfusion data was collected just before MCAO, immediately after initiation of ischemia, and immediately after intra-arterial injection. Average signal intensity was taken from fixed-size regions of interest drawn over the ipsi- and contralateral parietal bone plates. Ipsilateral CBF was reported as a percentage of contralateral CBF to indicate reperfusion efficacy.

### Intra-arterial (IA) injection procedure

To generate a successful IA injection procedure, microcatheters (Doccol, MA) and Hamilton microliter syringe (Hamilton, NV,100 µl) were used for injection. Mice were subjected to MCAO. At the beginning of reperfusion, the microcatheter was inserted carefully into internal carotid artery via the same surgical path used for occluding the MCA. The microcatheter was secured in place and the plunger on the Hamilton syringe was deployed so that 20 µl of methylene blue was incrementally administered over the course of 5 minutes. After administration, the microcatheter was kept in place for 2 minutes to mitigate backflow before being withdrawn and the surgical site was closed as described above. Mice were sacrificed, brains were extracted, and 2 mm serial sections were cut to display the distribution of dye in the cerebral vessels and parenchyma.

### IA DNase administration

The microcatheter was introduced to the internal carotid artery as described above. 25 µg/25µl DNase (n=10) or 25 µl normal saline (vehicle, n=10) were administered as described above. After administration, the microcatheter was withdrawn, the external carotid artery was permanently ligated, and the ICA and CCA were reopened. Post-operative care was performed as described in the above survival surgery. Cerebral blood flow was re-evaluated and recorded before mice were placed in the intensive care unit for recovery.

### Intraperitoneal (IP) DNase treatment

IP DNase administration: Mice were administered 50 µg/100 µl deoxyribonuclease I (DNase) (n=10) or normal saline (vehicle, n=10) via IP injection immediately after establishing reperfusion.

### 28-point sensorimotor neurological deficit score (NDS)

At 24 hours post-stroke, a 28-point rodent neurological deficit score (NDS) was used to evaluate sensorimotor function as described previously [27, 28]. The NDS uses a 0 (no deficits) to 4 (extreme deficits) point system on 7 sensorimotor functions: body symmetry, gait (open bench top), circling behavior, assessment of climbing, assessment of front limb symmetry, compulsory circling, and whisker response. To reduce bias in data collection, all mice, experimental groups, and treatments were randomly assigned and coded such that outcomes investigators were blinded. All testing was scored independently by two investigators who were blinded to the identity of the individual animals, and their scores were averaged.

### Wire hang test

The wire hang test assesses locomotor ability [29]. At 24 hours post-stroke, mice were brought to the wire and allowed to grasp the wire only with their two forepaws. The mice were released and allowed to hang for a maximum of 2 minutes. Latency to fall (sec) was recorded to evaluate grip strength and endurance.

### Triphenyltetrazolium chloride (TTC) Staining

At 24 hours post-stroke, mice were sacrificed and transcardially perfused with 0.01 M phosphate buffered saline (PBS) solution (pH 7.6). Whole brains were extracted and chilled on dry ice. Partially frozen brains were then sectioned into 2 mm coronal slices and incubated in 1.5% prewarmed 2,3,5-Triphenyltetrazolium chloride (TTC) solution. Slices were incubated until clear infarct boundaries were observed on both sides of each section. Sections were transferred into a 4% paraformaldehyde solution at 4 °C overnight and scanned the next day using an HP Scanjet 5590. Postfixed brain slices were cryoprotected with 30% sucrose in PBS overnight in preparation for serial coronal cryostat sections (16 µm).

### Assessment of ischemic infarct volume

An in-house image analysis program was written in Python approximating the infarct areas using a color filter to determine infarcted versus non-infarcted tissue to calculate the corresponding infarct volumes. The program determined each pixel’s red-green-blue color value (RGB values). The program then searched for and counted all pixels above a certain threshold of RGB value (closer to white). Since TTC stains healthy tissue red while infarcted tissue remains unstained (white), the program was able to accurately define infarct borders and count all the pixels within said border. To address variations in the color balance of images, the thresholds for these values can be adjusted by the investigator so that infarct volumes can be accurately calculated. The images were also filtered based on the size of the detected areas, allowing for removal of small non-infarct areas that were inaccurately detected by the color filter. The volume was then calculated based on the areas of the anterior and posterior sides, the thickness of the sections, and a pixels-to-mm^2^ conversion factor calculated from a reference point with a known area in mm^2^ (see equations below). The reference point used was a white plus sign that was present on every slide with a consistent area.

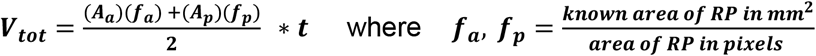

#### Calculations

the areas from the anterior and posterior faces of the sections were each converted to mm^2^ using their corresponding conversion factors (*f*_*a*,_ *f*_*p*_) and the two were averaged; the result was multiplied by the thickness of the sections to find the total volume of the infarct. *A*_*a*_ *= total area of anterior faces; A*_*p*_ *= total area of posterior faces; f*_*a*_ *= mm*^*2*^*/pixel factor from anterior view image; f*_*p*_ *= mm*^*2*^*/pixel factor from posterior view image; t = thickness of sections; RP = reference point of corresponding slide*.

### Immunohistochemistry staining, immunofluorescence staining, and imaging

Cryostat sections were thawed, washed, and rehydrated. For immunohistochemistry staining, sections were blocked with 3% animal serum in PBS-T for 1h and then incubated with anti-Ly6G (BD Bioscience, #551459) primary antibody overnight at 4°C. Sections were washed and incubated in anti-rabbit Horseradish peroxidase secondary antibody. Sections were then treated with DAB substrate working solution (Vector Laboratories, #SK-4100). Hematoxylin was used to counterstain cell nuclei. Images were collected using a Zeiss microscope and post-processed using ImageJ. For immunofluorescence staining, the sections were washed and rehydrated. Sections were incubated with the following primary antibodies: anti-Ly6G (BD Bioscience, #551459), anti-Vascular cell adhesion molecule 1 (VCAM1, Thermo Fisher, #MA5-11447), anti-myeloperoxidase (MPO, Bioss, BS-4943R), or Lectin - Dylight 488 (Vector, #DL-1174-1) overnight at 4°C. A secondary antibody (Alexa 488 or Alexa 568, Thermo Fisher) incubation was performed if necessary and DAPI counterstained cell nuclei. Images were collected using a Nikon confocal microscope and post-processed using Nikon Microscope Imaging Software and ImageJ.

### Microvascular density

Mice were subjected to MCAO as previously described. After reperfusion, mice were administered either 50ug/100ul DNase or normal saline solution through IP injection. Mice were sacrificed at 24 hours after reperfusion and transcardially perfused with PBS followed by 4% paraformaldehyde in PBS. This was followed by perfusion with 10 ml of 2% gelatin combined with albumin-FITC (Sigma, 0.125% w/v). The brain was extracted and incubated in 4% PFA overnight at 4°C. Postfixed brains were incubated in 15%, then 30% sucrose (pH 7.4). Serial coronal sections (16 µm) were prepared, and sections were imaged on Zeiss confocal laser microscope (LSM700, Carl Zeiss Microscopy).

### Quantification of NETs biomarkers concentration in plasma

Whole blood was centrifuged at 1500g for 15 minutes at 4°C for plasma collection. Plasma levels of cell-free DNA were measured by Quant-iT PicoGreen dsDNA Assay kit (ThermoFisher, #P11496) following the manufacturer’s protocol. Enzyme-linked immunosorbent assay (ELISA) kits were used to determine the plasma levels of citrullinated histone H3 (CitH3 (Clone 11D3) ELISA Kit, Cayman Chemicals, #501620), myeloperoxidase (Mouse MPO Quantikine ELISA kit, ThermoFisher, #EMMPO), and neutrophil elastase (Mouse Elastase Platinum ELISA, Abcam, #ab252356) according to each respective manufacturer’s protocol.

### Statistical analysis

All data are presented as mean ± standard error of mean. Unpaired t-test was used for data analysis from two groups and one-way ANOVA with post-hoc Tukey’s test was applied for the data from multiple groups. GraphPad Prism 10 (GraphPad, MA) was used for statistical analysis, DNase activity curve, and graphs generation. Statistical significance was defined as p < 0.05 for all analyses.

## 2. Results

### 2.1 Evaluation of Deoxyribonuclease I (DNase) activity in vitro and in vivo

To confirm the activity of Deoxyribonuclease I (DNase), our *in vitro* study showed that DNase exhibited a reliable dose-dependent activity curve (Fig 1.A). Our *in vivo* experiment detected the activity curve of serum DNase in mice 30 minutes after IV or IP DNase administration (Fig 1.B). The activity curve showed serum DNase activity was significantly elevated in both IP and IV groups in comparison to control serum. However, DNase via IV injection generated a quicker and stronger activity curve compared with DNase via IP injection. Endogenous DNase activity in vehicle controls only exhibited a linear activity curve (Fig 1B). This data indicates that DNase supplementation improves *in vivo* DNase activity, and that the method of administration has a significant effect on early DNase activity.

**Fig 1.**
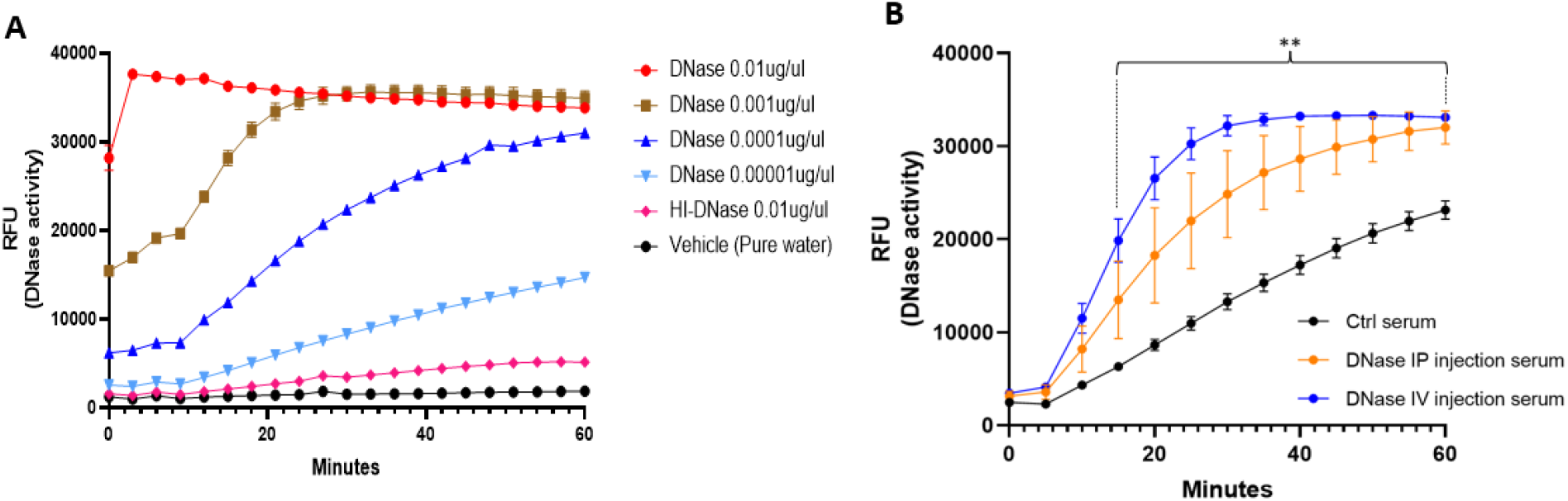
DNase activity in vitro and in vivo. **A)**. Dose dependent DNase activity curve in vitro. **B)**. DNase activity curve in mice serum. DNase IP injection, n=3; tail vein (IV) injection, n=4, and Ctrl, n=3. Note: ** p<0.01, both IP and TV injection serum vs. Ctrl serum. GraphPad Prism 9.5 was used for data and analysis and DNase activity curve generation.

### 2.2 Procedure of successful intra-arterial (IA) injection in mice with AIS

In this study, we focused on modeling a protocol that would have meaningful and immediate translation to current clinical practice for stroke therapy. Thus, we explored IA injection as a means of directly administering low-dose DNase to a local cerebral area. To build a successful IA injection procedure, microcatheters attached to a Hamilton microliter syringe were used to inject. We first utilized methylene blue to gage distribution via this method and as a proof-of-concept demonstration. Methylene blue was directly administered intra-arterially after MCAO and recanalization (Fig 2.A). Brain surface imaging clearly showed that the methylene blue was distributed along the micro vessels supplied by the middle cerebral artery in the dorsal and ventral views of the right hemisphere of the mouse brain (Fig 2.B). Coronal sections revealed that methylene blue was disseminated throughout the ischemic, but not contralateral hemisphere (Fig 2.C). Methylene blue was localized to the vasculature and parenchyma of the ischemic hemisphere but not the contralateral hemisphere. This indicates that methylene blue was successfully delivered via direct IA injection into the ischemic area of the mouse brain, and suggests that IA administration can successfully deliver local, low-dose therapeutics for LVOs.

**Fig 2.**
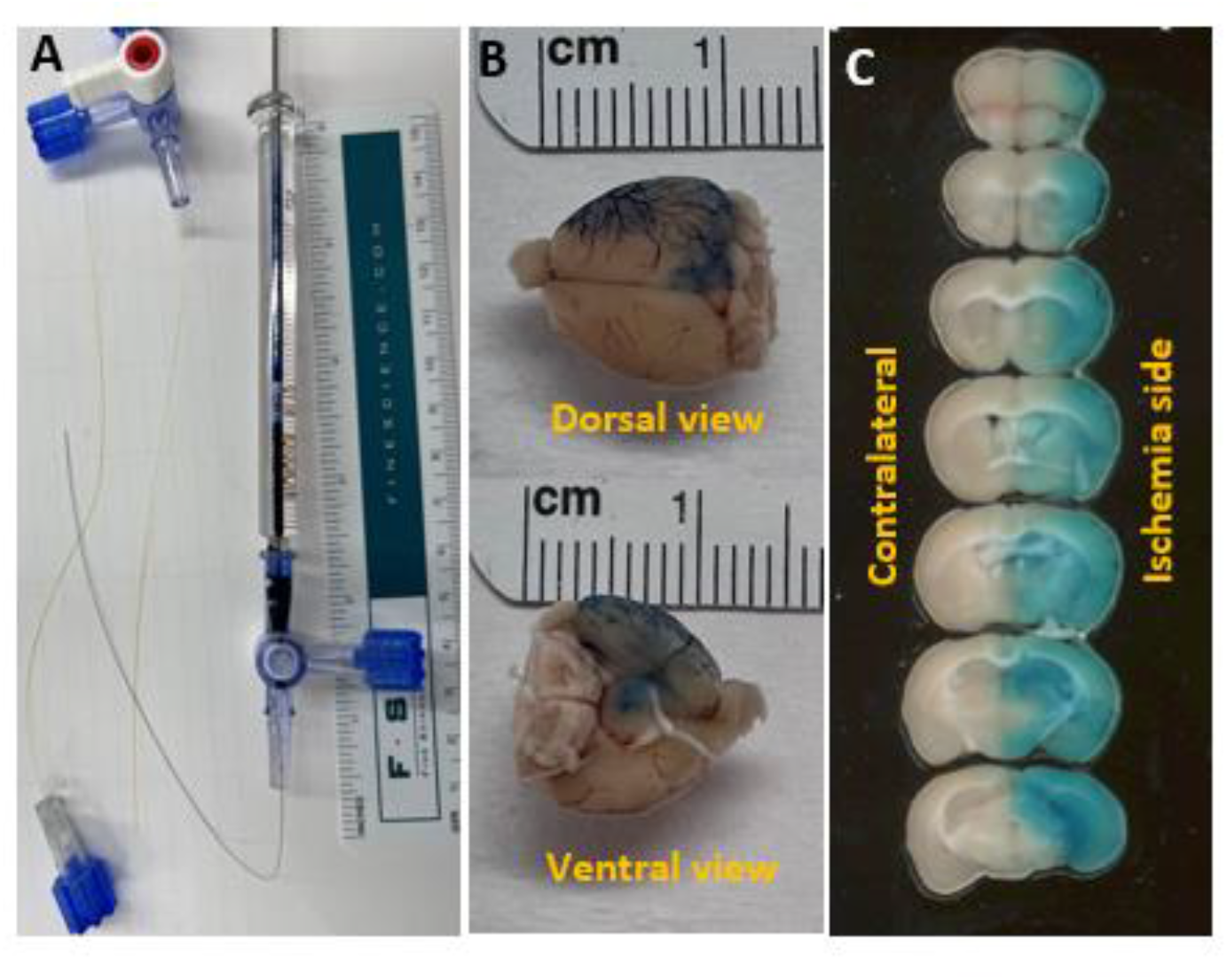
Intra-artery injection tools and Methylene blue injection after MCAO. A). Microcatheters and Hamilton syringe. B). Images of mouse brain after direct intra-artery Methylene blue injection, mouse was subjected to MCAO with 60’ ischemia followed by reperfusion and immediate intra-arterial 20 µl Methylene blue injection. C). Images of brain sections. With immediate intra-arterial methylene blue injection, blue dye spreads in ischemic reperfusion brain tissue side and no blue dye were observed in contralateral brain tissue.

### 2.3 IA DNase treatment improved neurological function and reduced infarct volume via increasing efficient cerebral reperfusion in aged mice with AIS

NETs contribute towards microcirculation hypoperfusion and are associated with worse prognosis [16-18, 20, 21]. Here, we are the first to investigate the effect of IA DNase administration in aged mice with AIS. In our study, laser imaging revealed that IA DNase injections post AIS improved reperfusion. The signal intensity ratios of cerebral blood flow images showed that IA DNase administration significantly improved CBF post-reperfusion in comparison to vehicle (Fig 3A, B). This emphasizes the potential of IA DNase administration in improving reperfusion efficiency in microvascular capillary beds post mechanical thrombectomy.

**Fig 3.**
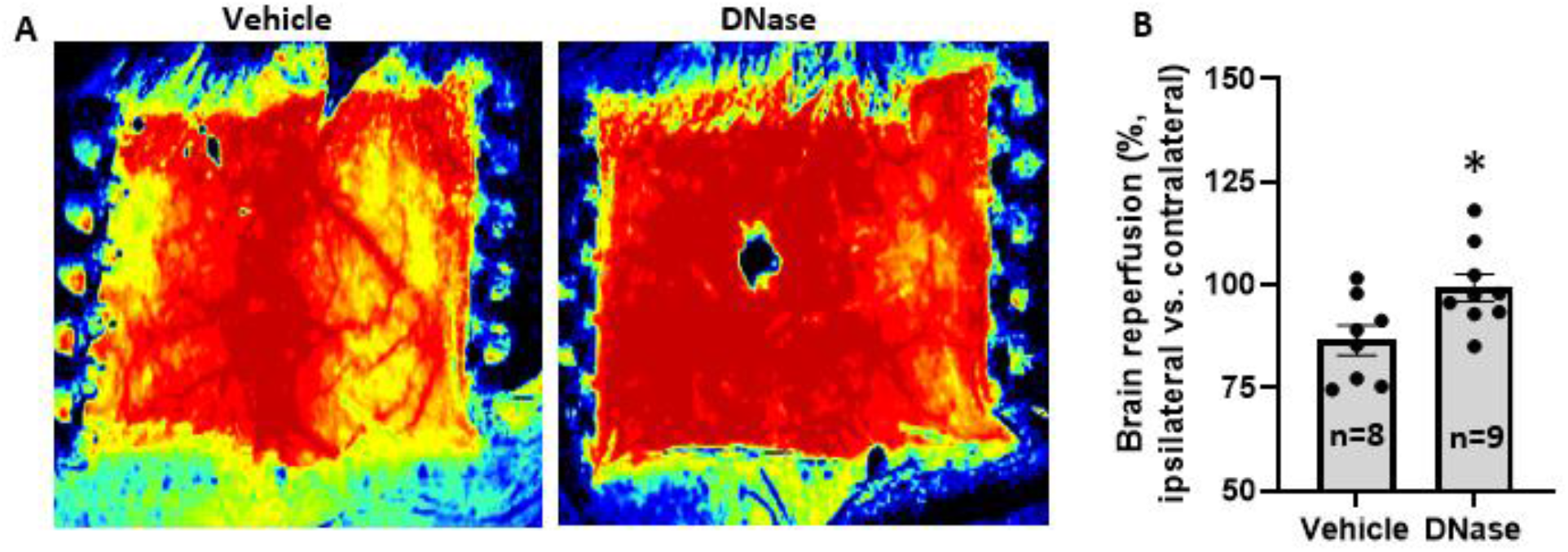
Intra-arterial DNase treatment improved early cerebral blood reperfusion in aged mice with AIS. Cerebral blood flow was evaluated immediately after intra-arterial DNase injection. The ratio of cerebral blood flow (ipsilateral (ischemic) side to contralateral side) was calculated. **A)**. Representatives of cerebral blood flow images for Vehicle and DNase, and **B)**. Results of the ratio of cerebral blood flow. Note: * p<0.05, compared with Vehicle.

This improved reperfusion efficiency corresponds with an improvement in functional neurological outcomes and a reduction in ischemic infarct volume. IA DNase administration significantly reduced neurological deficit scores and increased the latency to fall in wire hang test in comparison to those of mice receiving vehicle treatment (Fig 4A, B). These findings suggest that IA administration of DNase post-reperfusion improves sensorimotor outcomes in AIS mice. Additionally, IA DNase administration post-reperfusion produced an obvious reduction in stroke volume in comparison to vehicle controls (Fig. 4C). Volume quantifications confirmed a significant reduction in infarct volume in the IA DNase group in comparison to that of the vehicle controls (Fig. 4D). Taken together, these findings point to IA DNase administration post-recanalization improving stroke outcomes in AIS mice, possibly by mitigating futile reperfusion and improving cerebral blood flow.

**Fig 4.**
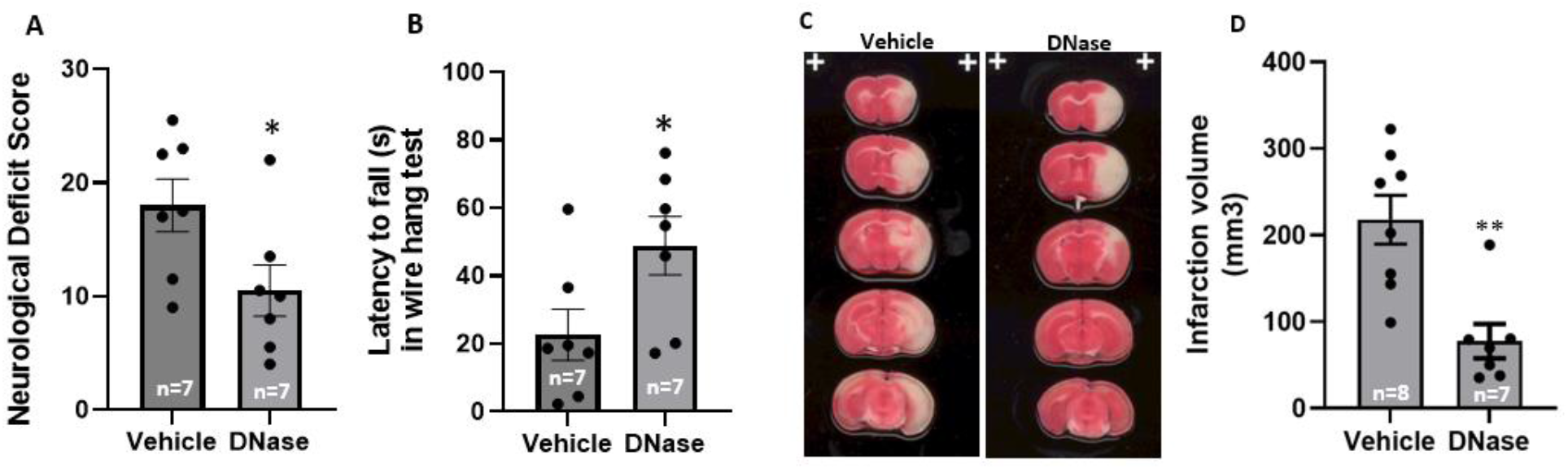
Intra-arterial DNase treatment improved neurological function outcomes and reduced ischemic infarction in aged mice with AIS. **A**) Neurological deficit score, **B**) Latency to fall in wire hang test. **C**) Representative TTC images, **D**) Bar graph of infarct volumes, python Image analysis program was used for infarct volume calculation based on TTC staining images. Note: * p<0.05, ** p <0.01, compared with Vehicle.

### 2.4 IA DNase reduced NETs formation in the penumbra of aged mice with AIS

To confirm the effect of IA DNase administration on NET formation, we evaluated NET deposition and neutrophil infiltration in both the parenchyma and micro vessels in the penumbra of mice with AIS. Fixed-frozen brain tissue was sectioned and stained with neutrophil and NET markers (Ly6G, MPO), as well as vascular markers (VCAM and Lectin) (Fig 5). The images show abundant strong neutrophil signals in the parenchyma and micro vessels in the penumbra of mice from vehicle treatment group (Fig 5. B-D, F-H). Conversely, DNase reduced the number of neutrophils significantly in the parenchyma and micro vessels in the penumbra of AIS mice (Fig 5. J-L, N-P). Statistical analysis indicated that DNase treatment significantly reduced the infiltrated neutrophils count in parenchyma compared to vehicle treatment (Fig 5.Q). These findings indicate that IA DNase administration reduces both the number of neutrophils and NETs in the penumbra of AIS mice in comparison to vehicle controls.

**Fig 5.**
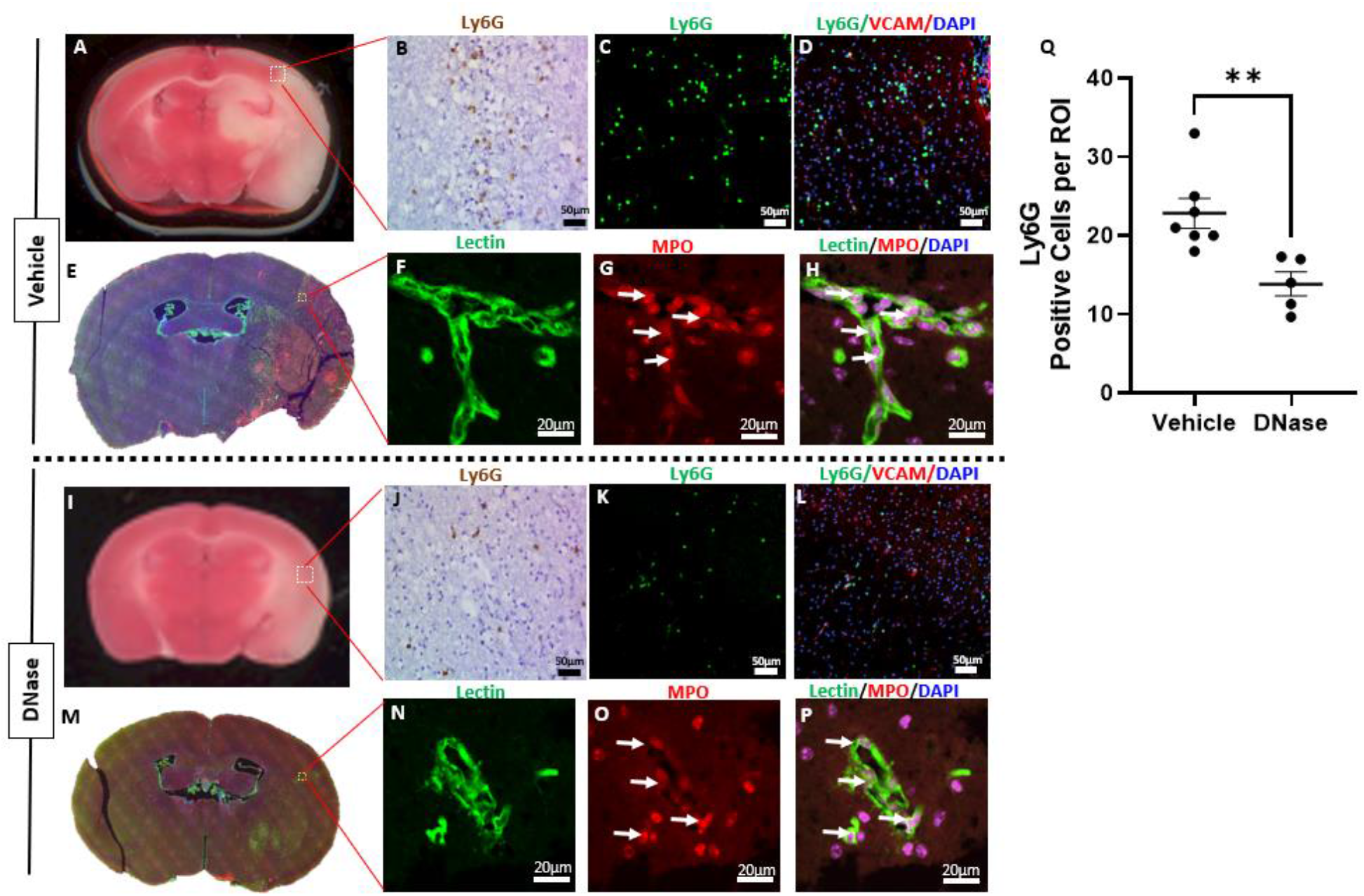
Intra-arterial DNase treatment reduced neutrophils infiltration and NETs formation in ischemic penumbra of aged mice with AIS. Brain sections were stained with neutrophil antibodies (Ly6G and myeloperoxidase-MPO), vascular marker antibodies (vascular cell adhesion molecule 1 (VCAM) and Lectin), and nuclei (DAPI). NETs were identified by Ly6G or MPO positive signaling. Representative images from vehicle treatment group: **A**) Image of TTC staining, penumbra area recircled with white dash line. **B**) Image of anti-Ly6G DAB staining. **C**) Image of anti-Ly6G immunofluorescence staining. **D**) Merge image of anti Ly6G (green), anti-VCAM (red), and DAPI (blue) immunofluorescence staining. **E**) Merge image of anti-Lectin (green), anti-MPO (red), and DAPI (blue) of whole section. **F**) Image of anti-Lectin. **G**) Image of anti-MPO, intra-vascular (white arrow). **H**) Merged image of anti-Lectin, anti-MPO, and DAPI, intra-vascular NETs signal (white arrow) in penumbra. Representative images from DNase treatment group: **I**) Image of TTC staining, penumbra area recircled with white dash line. **J**) Image of anti-Ly6G DAB staining. **K**) Image of anti-Ly6G immunofluorescence staining. **L**) Merged image of anti Ly6G (green), anti-VCAM (red), and DAPI (blue) immunofluorescence staining. **M**) Merged image of anti-Lectin (green), anti-MPO (red), and DAPI (blue) of whole section. **N**) Image of anti-Lectin. **O**) Image of anti-MPO, intra-vascular (white arrow). **P**) Merged image of anti-Lectin, anti-MPO, and DAPI, intra-vascular NETs signal (white arrow) in penumbra. **Q**) Plots graph of the number of anti-Ly6G positive signal. Note: ** p <0.01, compared with Vehicle.

### 2.5 IP DNase treatment reduced NETs formation in systemic circulation and improved capillary recanalization in the penumbra of mice with AIS

DNase treatment not only reduced NET burden in the ischemic penumbra, but also in systemic circulation. Our data showed that IP DNase treatment significantly reduced the plasma concentration of NE and dsDNA in mice with AIS when compared to that of the vehicle treatment group (Fig 6. A, B). NET containing cerebral microvascular thrombi may occlude capillary blood flow and induce penumbral hypoperfusion in stroke [15, 17, 20, 21]. To evaluate whether DNase treatment also improves microcirculation after recanalization, we tested the density of micro vessels in the ischemic penumbra of AIS aged mice. Our data showed that the microvascular density in the penumbra of DNase-treated, aged mice with AIS was significantly higher than that of vehicle-treated, aged mice with AIS (Fig 6. C, D). This data suggests that DNase improves perfusion in the ischemic penumbra, potentially by degrading intravascular NETs, particularly in micro vessels. Taken together, these findings indicate that DNase treatment reduces NET burden *in vivo* and provides a mechanistic explanation as to how DNase improves reperfusion and stroke outcomes in AIS.

**Fig 6.**
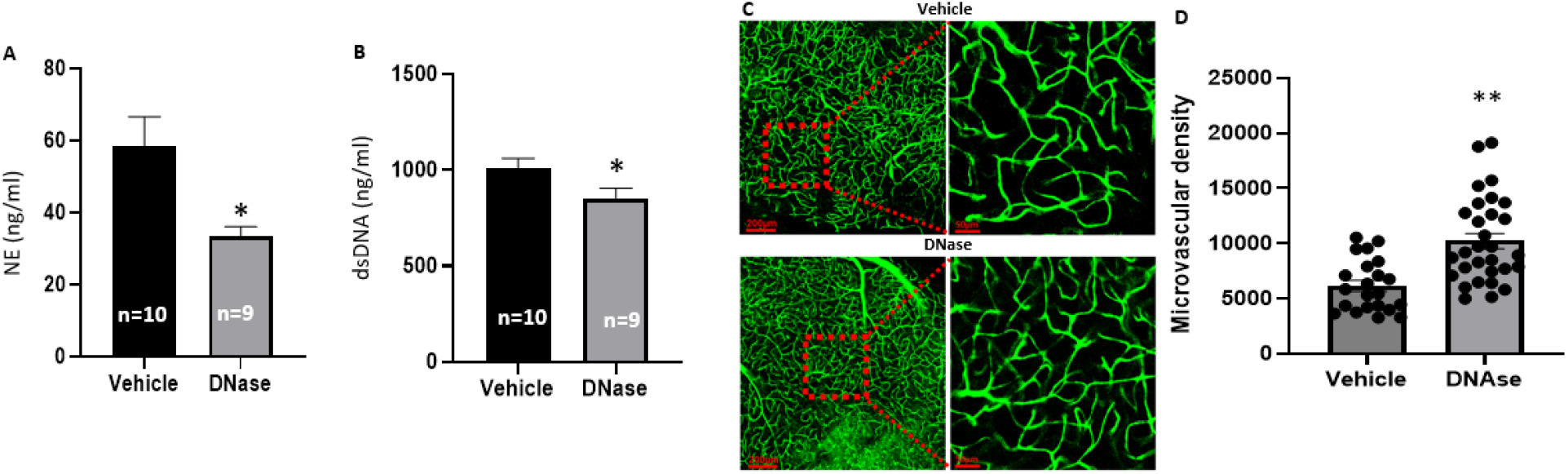
Intraperitoneal DNase treatment reduced NETs formation in systemic circulation and improved capillary recanalization in penumbra of mice with AIS. **A**) Plasma NE concentration. **B**) Plasma dsDNA concentration. **C**) Representative microvascular images taken from penumbra of mice with vehicle treatment (n=6) and DNase treatment (n=9). **D**) Microvascular density as calculated by fluorescent signal. Note: * p <0.05, ** p<0.01, compared with Vehicle.

## Discussion

Neutrophils are the first cell type to infiltrate ischemic tissue. Once there, they release web-like DNA structures (NETs) in the diseased brain parenchyma and cerebral microvasculature [30-33]. NET formation facilitates thrombosis and contributes to reperfusion resistance during the thrombolytic treatment of AIS [20, 34]. Additionally, plasma NET markers are significantly increased in ischemic stroke patients, correlating with worse outcomes, and accurately used to predict the stroke mortality and morbidity [16-18, 34-37]. Thus, approaches to reduce NET burden have beneficial potential in the treatment of AIS [15, 38-43].

DNase is FDA-approved for the treatment of cystic fibrosis [44, 45] and its safety and clinical efficacy in treating respiratory dysfunction is well established [46, 47]. However, there are no FDA approved therapies for modifying NETs with respect to stroke. Intravascularly, DNase breaks down NETs and other cell-free DNA, reducing blood viscosity and resolving vascular congestion [46]. Both intraperitoneal injection and tail vein injection of DNase therapy exhibited promising neuroprotective effects and improved neurological outcomes in young mice when given before and after acute ischemic stroke, suggesting the targeting of NETs with DNase as a promising therapeutic in AIS [15, 18, 19, 48, 49].

Our study of *in vitro* and *in vivo* DNase activity describes the comparative efficacy of tail vein versus intraperitoneal injection, which was previously unresolved. Not surprisingly, we found that the activity of DNase via tail vein administration was stronger and faster than that administrated through intraperitoneal injection. Importantly, this finding suggested to us that direct intra-arterial DNase administration may exhibit an even more robust response given its immediate application downstream of the occlusion site.

Therefore, we investigated direct intra-arterial administration as a new procedure in the delivery of stroke therapeutics and examined the effects of IA DNase administration in aged mice of both sexes with AIS. Like our own findings, a recent study reported that reperfusion was significantly impaired in aged mice following recanalization of MCA occlusion [50]. Additionally, they found vascular occlusions in the ischemic penumbra were associated with increased neutrophil infiltration and NETs. These phenomena were worse in aged mice, suggesting that the effects of relieving NET pathology could have a greater effect in aged mice than in young mice with AIS. Our direct intra-arterial DNase treatment increased cerebral blood flow and improved stroke outcomes in aged mice with AIS (Fig. 3,4). Additionally, IP DNase treatment prevented the reduction of capillary density and reduced plasma NET markers (NE and dsDNA) in aged mice with AIS (Fig. 5, 7). Taking both ours and others’ findings [50], it is hypothesized that the reduction of neutrophil infiltration and the disruption of microvascular NETs (Fig. 5, 6) reduces the formation of microvascular occlusions and may mitigate futile recanalization.

Futile recanalization (FR) continues to pose problems for physicians in the treatment of LVOs. Intravascular administration of tPA after mechanical thrombectomy has been proposed as a treatment for FR [51]. Reports describing mechanical thrombectomy in conjunction with tPA administration include improvement in stroke outcomes, whereas others describe increased occurrence of hemorrhagic transformation and intracerebral hemorrhage [52-54]. Interestingly, in addition to the disruption of NETs and providing neuroprotection in ischemic injury, DNase reduces tPA-associated hemorrhagic transformation by mitigating tPA-induced blood brain barrier breakdown and reducing tPA-associated hemorrhagic transformation [19, 55]. Direct intra-arterial DNase-tPA coadministration at the occlusion site may similarly reduce hemorrhagic transformation seen in mechanical thrombectomy with intra-arterial tPA treatment while maintaining its positive outcomes. Further study will be needed to test the safety of intra-arterial tPA-DNase coadministration after mechanical thrombectomy, and whether there is a clear clinical benefit to coadministration in comparison to intra-arterial DNase alone.

In summary, we explored direct intra-arterial DNase administration as a new procedure in stroke therapeutics in aged mice of both sexes with AIS. The direct intra-arterial DNase treatment significantly increased cerebral blood flow, improved neurological function, reduced cerebral infarct volume, and decreased NET burden in the penumbra of aged AIS mice. Our data demonstrate that direct intra-arterial injection of DNase at the site of occlusion successfully provides an immediate and efficient improvement of cerebral blood flow after recanalization in the treatment of LVOs. This study provides promising evidence in support of the clinical application of DNase treatments and IA administration.

## Reference

1. Collaborators GBDS: Global, regional, and national burden of stroke and its risk factors, 1990-2019: a systematic analysis for the Global Burden of Disease Study 2019. Lancet Neurol 2021, 20:795–820.

2. DALYs GBD, Collaborators H: Global, regional, and national disability-adjusted life-years (DALYs) for 359 diseases and injuries and healthy life expectancy (HALE) for 195 countries and territories, 1990-2017: a systematic analysis for the Global Burden of Disease Study 2017. Lancet 2018, 392:1859–1922.

3. Malhotra K, Gornbein J, Saver JL: Ischemic Strokes Due to Large-Vessel Occlusions Contribute Disproportionately to Stroke-Related Dependence and Death: A Review. Front Neurol 2017, 8:651.

4. Lattanzi S, Cuccurullo C, Orlandi N, Borzi G, Bigliardi G, Maffei S, Giovannini G, Meletti S: Futile recanalization is associated with increased risk of post-stroke epilepsy. J Neurol Sci 2024, 462:123067.

5. Schwarting J, Froelich MF, Kirschke JS, Mehrens D, Bodden J, Sepp D, Reis J, Dimitriadis K, Ricke J, Zimmer C, et al: Endovascular thrombectomy is cost-saving in patients with acute ischemic stroke with large infarct. Front Neurol 2024, 15:1324074.

6. Saber H, Desai SM, Haussen D, Al-Bayati A, Majidi S, Mocco J, Hassan AE, Rajah G, Waqas M, Davies JM, et al: Endovascular Therapy vs Medical Management for Patients With Acute Stroke With Medium Vessel Occlusion in the Anterior Circulation. JAMA Netw Open 2022, 5:e2238154.

7. Goyal M, Menon BK, van Zwam WH, Dippel DW, Mitchell PJ, Demchuk AM, Davalos A, Majoie CB, van der Lugt A, de Miquel MA, et al: Endovascular thrombectomy after large-vessel ischaemic stroke: a meta-analysis of individual patient data from five randomised trials. Lancet 2016, 387:1723–1731.

8. Shen H, Killingsworth MC, Bhaskar SMM: Comprehensive Meta-Analysis of Futile Recanalization in Acute Ischemic Stroke Patients Undergoing Endovascular Thrombectomy: Prevalence, Factors, and Clinical Outcomes. Life (Basel) 2023, 13.

9. Benjamin EJ, Blaha MJ, Chiuve SE, Cushman M, Das SR, Deo R, de Ferranti SD, Floyd J, Fornage M, Gillespie C, et al: Heart Disease and Stroke Statistics-2017 Update: A Report From the American Heart Association. Circulation 2017, 135:e146–e603.

10. Mijajlovic MD, Pavlovic A, Brainin M, Heiss WD, Quinn TJ, Ihle-Hansen HB, Hermann DM, Assayag EB, Richard E, Thiel A, et al: Post-stroke dementia - a comprehensive review. BMC Med 2017, 15:11.

11. Morishima I, Sone T, Okumura K, Tsuboi H, Kondo J, Mukawa H, Matsui H, Toki Y, Ito T, Hayakawa T: Angiographic no-reflow phenomenon as a predictor of adverse long-term outcome in patients treated with percutaneous transluminal coronary angioplasty for first acute myocardial infarction. J Am Coll Cardiol 2000, 36:1202–1209.

12. Kitano T, Todo K, Yoshimura S, Uchida K, Yamagami H, Sakai N, Sakaguchi M, Nakamura H, Kishima H, Mochizuki H, et al: Futile complete recanalization: patients characteristics and its time course. Sci Rep 2020, 10:4973.

13. Baek JH, Kim BM, Kang DH, Heo JH, Nam HS, Kim YD, Hwang YH, Kim YW, Kim YS, Kim DJ, et al: Balloon Guide Catheter Is Beneficial in Endovascular Treatment Regardless of Mechanical Recanalization Modality. Stroke 2019, 50:1490–1496.

14. Niccoli G, Burzotta F, Galiuto L, Crea F: Myocardial no-reflow in humans. J Am Coll Cardiol 2009, 54:281–292.

15. Laridan E, Denorme F, Desender L, Francois O, Andersson T, Deckmyn H, Vanhoorelbeke K, De Meyer SF: Neutrophil extracellular traps in ischemic stroke thrombi. Ann Neurol 2017, 82:223–232.

16. Valles J, Lago A, Santos MT, Latorre AM, Tembl JI, Salom JB, Nieves C, Moscardo A: Neutrophil extracellular traps are increased in patients with acute ischemic stroke: prognostic significance. Thromb Haemost 2017, 117:1919–1929.

17. Demyanets S, Stojkovic S, Mauracher LM, Kopp CW, Wojta J, Thaler J, Panzer S, Gremmel T: Surrogate Markers of Neutrophil Extracellular Trap Formation are Associated with Ischemic Outcomes and Platelet Activation after Peripheral Angioplasty and Stenting. J Clin Med 2020, 9.

18. Denorme F, Portier I, Rustad JL, Cody MJ, de Araujo CV, Hoki C, Alexander MD, Grandhi R, Dyer MR, Neal MD, et al: Neutrophil extracellular traps regulate ischemic stroke brain injury. J Clin Invest 2022, 132.

19. Kang L, Yu H, Yang X, Zhu Y, Bai X, Wang R, Cao Y, Xu H, Luo H, Lu L, et al: Neutrophil extracellular traps released by neutrophils impair revascularization and vascular remodeling after stroke. Nat Commun 2020, 11:2488.

20. Kim SW, Lee JK: Role of HMGB1 in the Interplay between NETosis and Thrombosis in Ischemic Stroke: A Review. Cells 2020, 9.

21. Ducroux C, Di Meglio L, Loyau S, Delbosc S, Boisseau W, Deschildre C, Ben Maacha M, Blanc R, Redjem H, Ciccio G, et al: Thrombus Neutrophil Extracellular Traps Content Impair tPA-Induced Thrombolysis in Acute Ischemic Stroke. Stroke 2018, 49:754–757.

22. Mengozzi L, Barison I, Maly M, Lorenzoni G, Fedrigo M, Castellani C, Gregori D, Maly P, Matej R, Tousek P, et al: Neutrophil Extracellular Traps and Thrombolysis Resistance: New Insights for Targeting Therapies. Stroke 2024, 55:963–971.

23. Hendrix P, Witsch J, Spalart V, Schneider H, Oertel J, Geisel J, Martinod K, Hemmer S: Neutrophil extracellular trap biomarkers in aneurysmal subarachnoid hemorrhage: early decline of DNase 1 activity associated with delayed cerebral ischemia. Front Neurol 2024, 15:1354224.

24. Zeineddine HA, Hong SH, Peesh P, Dienel A, Torres K, Thankamani Pandit P, Matsumura K, Huang S, Li W, Chauhan A, et al: Neutrophils and Neutrophil Extracellular Traps Cause Vascular Occlusion and Delayed Cerebral Ischemia After Subarachnoid Hemorrhage in Mice. Arterioscler Thromb Vasc Biol 2024, 44:635–652.

25. Carminita E, Crescence L, Brouilly N, Altie A, Panicot-Dubois L, Dubois C: DNAse-dependent, NET-independent pathway of thrombus formation in vivo. Proc Natl Acad Sci U S A 2021, 118.

26. Yin J, Han P, Tang Z, Liu Q, Shi J: Sirtuin 3 mediates neuroprotection of ketones against ischemic stroke. J Cereb Blood Flow Metab 2015, 35:1783–1789.

27. Clark WM, Lessov NS, Dixon MP, Eckenstein F: Monofilament intraluminal middle cerebral artery occlusion in the mouse. Neurol Res 1997, 19:641–648.

28. Clark W, Gunion-Rinker L, Lessov N, Hazel K: Citicoline treatment for experimental intracerebral hemorrhage in mice. Stroke 1998, 29:2136–2140.

29. Ji S, Kronenberg G, Balkaya M, Farber K, Gertz K, Kettenmann H, Endres M: Acute neuroprotection by pioglitazone after mild brain ischemia without effect on long-term outcome. Exp Neurol 2009, 216:321–328.

30. Brinkmann V, Reichard U, Goosmann C, Fauler B, Uhlemann Y, Weiss DS, Weinrauch Y, Zychlinsky A: Neutrophil extracellular traps kill bacteria. Science 2004, 303:1532–1535.

31. Yuen J, Pluthero FG, Douda DN, Riedl M, Cherry A, Ulanova M, Kahr WH, Palaniyar N, Licht C: NETosing Neutrophils Activate Complement Both on Their Own NETs and Bacteria via Alternative and Non-alternative Pathways. Front Immunol 2016, 7:137.

32. Sollberger G, Tilley DO, Zychlinsky A: Neutrophil Extracellular Traps: The Biology of Chromatin Externalization. Dev Cell 2018, 44:542–553.

33. Yousefi S, Stojkov D, Germic N, Simon D, Wang X, Benarafa C, Simon HU: Untangling “NETosis” from NETs. Eur J Immunol 2019, 49:221–227.

34. Novotny J, Oberdieck P, Titova A, Pelisek J, Chandraratne S, Nicol P, Hapfelmeier A, Joner M, Maegdefessel L, Poppert H, et al: Thrombus NET content is associated with clinical outcome in stroke and myocardial infarction. Neurology 2020, 94:e2346–e2360.

35. Rainer TH, Wong LK, Lam W, Yuen E, Lam NY, Metreweli C, Lo YM: Prognostic use of circulating plasma nucleic acid concentrations in patients with acute stroke. Clin Chem 2003, 49:562–569.

36. Bang OY, Chung JW, Cho YH, Oh MJ, Seo WK, Kim GM, Ahn MJ: Circulating DNAs, a Marker of Neutrophil Extracellular Traposis and Cancer-Related Stroke: The OASIS-Cancer Study. Stroke 2019, 50:2944–2947.

37. Tsai NW, Lin TK, Chen SD, Chang WN, Wang HC, Yang TM, Lin YJ, Jan CR, Huang CR, Liou CW, Lu CH: The value of serial plasma nuclear and mitochondrial DNA levels in patients with acute ischemic stroke. Clin Chim Acta 2011, 412:476–479.

38. Hamam HJ, Khan MA, Palaniyar N: Histone Acetylation Promotes Neutrophil Extracellular Trap Formation. Biomolecules 2019, 9.

39. Neeli I, Radic M: Opposition between PKC isoforms regulates histone deimination and neutrophil extracellular chromatin release. Front Immunol 2013, 4:38.

40. Li P, Li M, Lindberg MR, Kennett MJ, Xiong N, Wang Y: PAD4 is essential for antibacterial innate immunity mediated by neutrophil extracellular traps. J Exp Med 2010, 207:1853–1862.

41. Wang Y, Li M, Stadler S, Correll S, Li P, Wang D, Hayama R, Leonelli L, Han H, Grigoryev SA, et al: Histone hypercitrullination mediates chromatin decondensation and neutrophil extracellular trap formation. J Cell Biol 2009, 184:205–213.

42. Neeli I, Khan SN, Radic M: Histone deimination as a response to inflammatory stimuli in neutrophils. J Immunol 2008, 180:1895–1902.

43. Martinod K, Demers M, Fuchs TA, Wong SL, Brill A, Gallant M, Hu J, Wang Y, Wagner DD: Neutrophil histone modification by peptidylarginine deiminase 4 is critical for deep vein thrombosis in mice. Proc Natl Acad Sci U S A 2013, 110:8674–8679.

44. Suri R: The use of human deoxyribonuclease (rhDNase) in the management of cystic fibrosis. BioDrugs 2005, 19:135–144.

45. Wagener JS, Kupfer O: Dornase alfa (Pulmozyme). Current Opinion in Pulmonary Medicine 2012, 18:609–614.

46. Fuchs HJ, Borowitz DS, Christiansen DH, Morris EM, Nash ML, Ramsey BW, Rosenstein BJ, Smith AL, Wohl ME: Effect of aerosolized recombinant human DNase on exacerbations of respiratory symptoms and on pulmonary function in patients with cystic fibrosis. The Pulmozyme Study Group. N Engl J Med 1994, 331:637–642.

47. Quan JM, Tiddens HA, Sy JP, McKenzie SG, Montgomery MD, Robinson PJ, Wohl ME, Konstan MW: A two-year randomized, placebo-controlled trial of dornase alfa in young patients with cystic fibrosis with mild lung function abnormalities. J Pediatr 2001, 139:813–820.

48. De Meyer SF, Suidan GL, Fuchs TA, Monestier M, Wagner DD: Extracellular chromatin is an important mediator of ischemic stroke in mice. Arterioscler Thromb Vasc Biol 2012, 32:1884–1891.

49. Martinod K, Wagner DD: Thrombosis: tangled up in NETs. Blood 2014, 123:2768–2776.

50. Gullotta GS, De Feo D, Friebel E, Semerano A, Scotti GM, Bergamaschi A, Butti E, Brambilla E, Genchi A, Capotondo A, et al: Age-induced alterations of granulopoiesis generate atypical neutrophils that aggravate stroke pathology. Nat Immunol 2023, 24:925–940.

51. Deng G, Chu YH, Xiao J, Shang K, Zhou LQ, Qin C, Tian DS: Risk Factors, Pathophysiologic Mechanisms, and Potential Treatment Strategies of Futile Recanalization after Endovascular Therapy in Acute Ischemic Stroke. Aging Dis 2023, 14:2096–2112.

52. Renu A, Millan M, San Roman L, Blasco J, Marti-Fabregas J, Terceno M, Amaro S, Serena J, Urra X, Laredo C, et al: Effect of Intra-arterial Alteplase vs Placebo Following Successful Thrombectomy on Functional Outcomes in Patients With Large Vessel Occlusion Acute Ischemic Stroke: The CHOICE Randomized Clinical Trial. JAMA 2022, 327:826–835.

53. Suzuki K, Matsumaru Y, Takeuchi M, Morimoto M, Kanazawa R, Takayama Y, Kamiya Y, Shigeta K, Okubo S, Hayakawa M, et al: Effect of Mechanical Thrombectomy Without vs With Intravenous Thrombolysis on Functional Outcome Among Patients With Acute Ischemic Stroke: The SKIP Randomized Clinical Trial. JAMA 2021, 325:244–253.

54. Desilles JP, Loyau S, Syvannarath V, Gonzalez-Valcarcel J, Cantier M, Louedec L, Lapergue B, Amarenco P, Ajzenberg N, Jandrot-Perrus M, et al: Alteplase Reduces Downstream Microvascular Thrombosis and Improves the Benefit of Large Artery Recanalization in Stroke. Stroke 2015, 46:3241–3248.

55. Wang R, Zhu Y, Liu Z, Chang L, Bai X, Kang L, Cao Y, Yang X, Yu H, Shi MJ, et al: Neutrophil extracellular traps promote tPA-induced brain hemorrhage via cGAS in mice with stroke. Blood 2021, 138:91–103.

